# Use of Bioelectric Impedance Analysis (BIA) as a new method to detect prostate cancer

**DOI:** 10.1101/2020.02.12.943829

**Authors:** Riccardo Bartoletti, Alberto Greco, Tommaso Di Vico, Jacopo Durante, Vincenzo Ficarra, Enzo Pasquale Scilingo, Gaetano Valenza

**Author notes:** **Corresponding author**: Prof. Riccardo Bartoletti, University Urology Unit, Cisanello Hospital Bld. 30 C, Via Paradisa 4, 56124 PISA, Italy, Mailbox.

## Abstract

**Background:** To determine the accuracy of a novel BIA test endorectal probe.

**Methods:** One hundred-forty consecutive patient candidates to prostate biopsy and 40 healthy volunteers were selected (NCT03428087). Total PSA and PSA density (PSAD) determinations, digital rectal examination (DRE), and the BIA test were analysed in patients and controls. A 16 cores trans rectal prostate biopsy was performed on all patients with clinical suspicion of PCa after a multiparametric MRI (mMRI) test. The study endpoints were to determine accuracy of BIA test in comparison to PSA, PSAD levels, and mMRI and obtain PCa prediction in candidates to prostate biopsy by BIA test. The Mann-Withney U test, the Wilkoxon rank test, and Holm-Bonferroni’s method were adopted for statistical analyses, and a computational approach was also applied to differentiate patients with PCa from those with benign disease (BPH).

**Results:** Combined DRE, TRUS, PSA, and PSAD alone failed to satisfactorily discern patients with PCa from those with BPH (62.86% of discrimination accuracy) and mMRI PIRADS ≥3 showed a sensitivity of 83% and a specificity of 59%. The accuracy in discerning PCa and BPH increased up to 75% by BIA test (sensitivity 63.33% and specificity 83.75%).

**Conclusions:** The BIA test is a simple, promising, cheap, and reliable test for PCa non-invasive diagnosis. The novel finger probe may improve PCa detection also in patients with low-risk PCa, thus reducing the need of useless biopsies.

## Introduction

Prostate cancer (PCa) diagnosis necessarily implies the use of prostate biopsy, which is an invasive procedure burdened by potentially relevant complications such as bleeding and systemic infection. On the other hand, currently available non-invasive diagnostic tools seem to be unable to reduce the number of unnecessary biopsies. The decision-making process for prostate biopsy is mainly based on total Prostate Specific Antigen (tPSA) values and the results of multiparametric MRI (mMRI). PSA levels alone are often unable to differentiate PCa from other benign prostate diseases with a sensitivity in PCa detection close to 32% [1]. The combination of total PSA and digital rectal examination (DRE) improve the PCa detection as well as the combination of PSA, DRE and trans-rectal ultrasound (TRUS) with a cancer detection rate of 50% [1,2]. mMRI of prostate in naïve patients with suspicion of prostate cancer remains of difficult application due to the high costs of the procedure and the high number of men who need to be investigated in every day clinical practice, although its accuracy has been improved by the PIRADS V2 score classification [3,4]. Therefore, there is a need for alternative non-invasive tools improving the selection of patient candidate for prostate biopsies.

Previous studies on BIA revealed enthusiastic results mainly in patients with aggressive cancers [5–7]. Studies conducted in the setting of prostate cancer in particular, were limited by the applicability of proper probes on the prostate gland surface and by the anatomic deep site in which the prostate is placed inside the pelvic bone girdle [8]. With the aim to reduce previous limitations in the applications of BIA test, we tested the performance of a novel endorectal probe in a series of consecutive patient candidate to prostate biopsy for suspicious prostate cancer.

The objectives of the present study were: 1) to test the accuracy of BIA test to detect prostate cancer; 2) to evaluate if the proposed bioimpedence-based methodology has to be further optimized to obtain clinically meaningful results in terms of reliability and repeatability; and 3) to develop a multi-feature decision support system including bioimpedence-based parameters for the prediction of prostate cancer.

## Patients and Methods

### Patient selection

All the patients who were candidates for a prostate biopsy for suspicious prostate cancer were consecutively and prospectively selected in the timeframe between March and September 2018. Presumptive diagnosis was based on persistently raised total PSA value (> 4 ng/mL) and/or suspicion of cancer at DRE. Patients younger than 45 years, those with a BMI higher than 28, and those affected by other neoplasms, diabetes, electrolyte imbalance, liver diseases, over hydration or dehydration were excluded from the study. A total of 123 patients with persistently high total PSA levels and negative DRE underwent to mMRI and were classified in accordance with the PIRADS V2 system. [4]. Moreover, a group of young healthy volunteers were collected from a series of patient candidates to circumcision and selected with the same exclusion criteria. Healthy voluntaries were enrolled if total PSA value was <4 ng/ml and no earlier history of prostate diseases or prostate-related symptoms were referred. The study protocol was developed in accordance with the Prisma Statement and approved by Internal Review Committee (IRC) (1251/2017) and then registered (NCT03428087). All patients and healthy volunteers provided their preliminary approval to participate in the study by signing an informed consent form.

For every patient the following pre-bioptical parameters were collected: age, BMI, baseline total PSA (ng/mL), digital rectal examination (DRE), prostate volume estimated during TRUS examination, PSA density and PIRADS score when available.

### The BIA test and the new endo rectal probe

Phase sensitive instruments are able to simultaneously measure Reactance, Resistance and provide the Phase Angle degree. Very low Phase angle values indicate cells with altered electrical activity due to different intracellular content, DNA and water in cancers.[9].

All patients underwent a BIA test using a new endorectal “finger probe” before to perform prostate biopsy. The Akern’s BIA tester is tested and validated instrument and was previously used to measure the biometric parameters [10]. The BIA test was provided with the patient placed in a left flank position as normally adopted for DRE and prostate biopsy procedures. The negative pole electrodes were placed at the base of penile shaft and at the coccyx apex while the positive ones at the penis tip and at the novel “finger probe” respectively **(Fig.1).** The electrodes placement was done to create a restricted electric field in the prostate area and get more reliable results in terms of sensitivity as demonstrated by other authors. [7,8,11,12]. The novel probe was conceived to test the prostate tissue directly and consists of an electrode placement over a single rubber finger glove tip wearable over the rubber gloves normally used to detect prostate abnormalities. Carbon fibers are fixed at the tip of the probe, passed into the rubber finger and connected to the “receptor” positive pole BIA electrode. **(Fig.1**). The use of flat and tender fibers other than rigid sensors, was planned to allow an easy and sensitive concomitant palpation of the prostate gland. Different registrations have been made for the two prostate lobes. The BIA tester automatically calculated resistance, Reactance and Phase angle.

**Fig.1.**
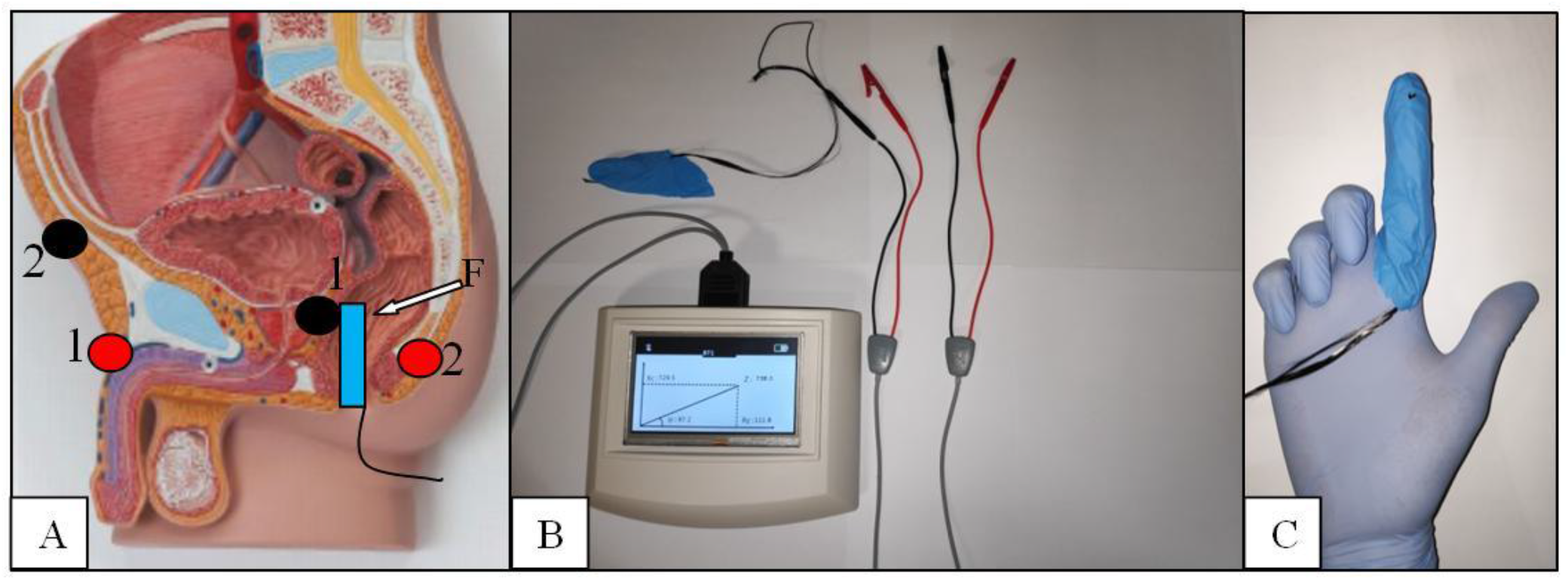
**A)** BIA electrodes placement during the BIA test: the negative pole electrodes are placed at the penis base and under the coccyx apex to avoid electrical interferences with pelvic bones. Electric field receptors (positive pole electrodes) are placed just up the pubic bone and on the tip of the finger probe (F). Each black electrode is connected to a proper electric field generator electrode (red) 1-1 and 2-2. **B)** The The Akern’s BIA tester. **C)** The “finger” probe: carbon fibers are placed on the tip of a second finger rubber glove tip and connected to the black BIA electrode through a small electro conducible metallic clamp.

### Prostate biopsy

Transrectal prostate biopsy was performed after antibiotic prophylaxis and local anaesthesia. In all cases at least 16 cores were systematically taken with systematic criteria except for the patients with PIRADS V2 score ≥3 who received adjunctive cores in relation to the number of prostate gland sites described at mMRI.[4]. Prostatic cores were embedded in formalin solution and then analyzed by two uro-pathologists.

### Statistical analysis, ROC curve, and SVM classification

All BIA measures were normalized by dividing their value by the prostate volume, which was estimated during the TRUS examination. Therefore, the BIA test was not dependent by the volume of the prostate, which can be significantly affected by the presence of cancer.

Continuous variables were statistically described using median and median absolute deviation (MAD) given the non-normality of the majority of samples demonstrated using Kolmogorov-Smirnov tests [13]. Accordingly, the Mann–Whitney U-test was used to statistically compare continuous variables from two different groups (e.g., patients vs. controls), whereas the Wilcoxon signed-rank test was used to compare differences in paired data [14,15]. To this extent, we statistically compared the bioimpedance features [resistance (RES), reactance (REA), and phase angle (PHASE)] estimated from the right and left prostate lobes in patients with diagnosed cancer data.

All p-values were corrected for multiple comparisons following the Holm-Bonferroni’s method [16].

A Receiver operating characteristic (ROC) curve analysis was performed on PIRADS V2 scores gathered from 123 patients by pairing false positive rates (1-specificity) and true positive rates (sensitivity) at different PIRADS V2 score thresholds [4,17]

To maximize a direct clinical applicability of the proposed study and move to a clinical evaluation at a single-patient level, we implemented a multi-feature computational approach that takes into account all features, combine them through a particular mathematical function, and automatically estimate the multi-dimensional threshold to be used to make a clinical decision for the prediction of cancer presence. The computational methodology is quite common in the bioengineering field and is named Support Vector Machine (SVM). We further extend the implementation of such a decision support system by integrating the so-called Recursive Feature Elimination (RFE) approach. This scores each patient’s feature such that it is possible to rank and select the most informative clinical information for the automatic discrimination of patients with prostate cancer and Benign Prostate Hyperplasia (BPH).

The prostate cancer recognition accuracy, as well as sensitivity and specificity of the proposed decision support system has been evaluated by implementing the so-called leave-one-subject-out procedure (LOSO), through which, considering N subjects, iteratively we split the clinical feature-set in a training set, comprising data from (N−1) patients, and a test set comprising data from the remaining one.

## Results

One hundred-forty men candidate to prostate biopsy for clinical suspicion of PCa were enrolled during the study period. No patients had relevant complication after the biopsy except for persistent one-month bleeding in seminal fluid. Patients with total PSA levels <4 ng/ml. presented suspicion of cancer at DRE and 6 out 15 PIRADS ≥3 at mMRI. Cancer was detected in 4 out 15 cases. PIRADS score ≥3 was found in 3 out 4 subjects with PCa. Cancer was found in 31 out 64 patients with total PSA between 4.1 and 10 ng/ml. PIRADS score ≥3 was found in 34 out 58 cases but only 22 of them presented association with PCa. Similarly, PCa was found in 21 out 61 patients with total PSA >10 ng/ml. MMRI confirmed the presence of cancer in 18/25 patients although resulted positive in 31 out 54 subjects. In 60 (42.8%) cases (median BMI 26.25 IQR 24.87 - 28.7) the biopsies resulted positive for prostate cancer while in the remaining 80 (57.2%) cases (median BMI 25.75 IQR 24.17 - 27.87) a non-neoplastic prostatic condition (BPH or inflammation) was diagnosed. ROC curve analysis performed on PIRADS V2 scores obtained from 123 mMRI of patients underwent to prostate core biopsy (no healthy volunteers have been included), showed for score threshold major or equal 3 a sensitivity of 83% and a specificity of 59%, VPP and VPN were 61% and 82% respectively.

By analyzing patients with prostate cancer, in 21 (35%) cases the disease involved a single lobe (11 the right side and 10 the left side). Conversely, in the other 39 (65%) cases both lobes were involved by the tumor. Therefore, according to D’Amico risk classification, 31 (51.6%) patients were classified as low-risk, 9 (15%) as intermediate and 20 (33.4%) as high-risk. The forty young healthy volunteers showed a median age of 37 (MAD=4) years and a median BMI of 25 (MAD=1.1). The median prostate volume was 19.23 cc (MAD=6.5) with a median PSA density of 0.05 (MAD=0.03). Comprehensive descriptive statistics of patients’ characteristics stratified according to the biopsy results are reported in **Table 1**.

**Table 1:** Patients’ characteristics. Descriptive ranges are expressed as (Median ± MAD). MAD = Median Absolute Deviation.

|  | PCa | BPH | Healthy<br>volunteers |
| --- | --- | --- | --- |
| Patients (n.) | 60 | 80 | 35 |
| Age (years) | 70 ± 5 | 69 ± 5 | 37 ± 4 |
| BMI (Kg/m<sup>2</sup>) | 26.01 ± 1.51 | 26.15 ± 2.15 | 25 ± 1.10 |
| Pre-biopsy total PSA value (ng/ml) |  |  |  |
| <4 | 4 | 11 | 35 |
| 4-10 | 31 | 33 | 0 |
| >10 | 25 | 36 | 0 |
| Prostate nodule/s at DRE (n.pts) |  |  |  |
| Right | 18 | 14 | 0 |
| Left | 10 | 7 | 0 |
| Bilateral | 8 | 4 | 0 |
| Negative | 24 | 55 | 35 |
| Prostate Volume at TRUS (ml) | 41.48 ± 11.88 | 53.78 ± 13.30 | 19.23 ± 6.50 |
| PSA Density (PSA/Volume) | 0.25 ± 0.13 | 0.17 ± 0.07 | 0.05 ± 0.03 |
| Prostate biopsy | YES | YES | NO |
| Low risk PCa (n.) | 31 | - | - |
| Intermediate risk PCa (n.) | 9 | - | - |
| High risk PCa (n.) | 20 | - | - |
| Patients who underwent to pre-biopsy mMRI (n.) | 60 | 63 | - |
| Patients with mMRI PIRADS≥3 (n./%) | 43/71.6 | 28/44.4 | - |
| Patients with mMRI PIRADS≥4 (n./%) | 15/25 | 2/3.1 | - |

Inferential statistics between patients with PCs, BPH, and controls are reported in **Table 2**. Comparing patients with PCa vs. controls, differences in age, PSA, prostate volume, and PSA density were found. Same statistical differences were found comparing BPH vs. controls (see **Table 2**). Concerning BPH vs. patients with PCa, significant differences were found in the prostate volume exclusively (p<0.01). Due to this reason PSAD analysis was introduced in the methodological accuracy determination to obtain normalized values.

**Table 2:**
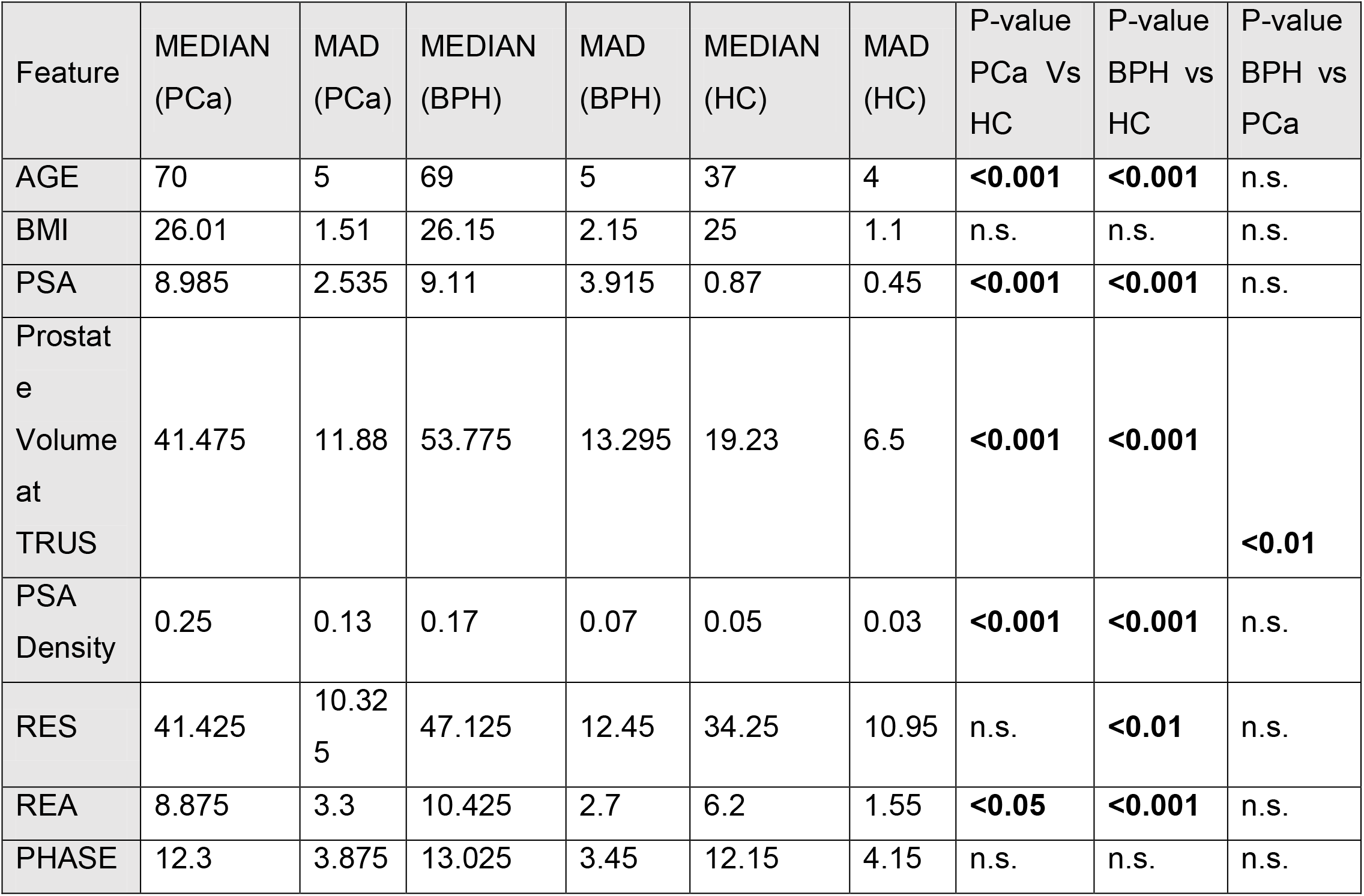
Statistical comparison between patients who underwent to prostate biopsy (cancer and BPH) and controls (n.s.: non-significant p-value). MAD = Median Absolute Deviation. BMI= Body Mass Index PSA= Prostate Specific Antigen TRUS= Trans Rectal Ultra Sound RES= Resistance REA= Reactance PHASE= Phase angle PCa = prostate cancer BPH= benign prostate hyperplasia HC= healthy controls

### BIA test parameters

**Table 2** summarizes and compares BIA test parameters gathered from patients with prostate cancer, benign prostatic disease, and healthy volunteers. While no significant differences between groups were found on the BIA parameter PHASE, significant differences were found in comparing BPH vs. controls using RES (p<0.01), as well as comparing controls vs. patients with PCa and controls vs. BPH using REA (p<0.05).

Concerning the statistical comparison between the three bioimpedance test measurements from the right and left side of the prostate, we split the dataset in three subsets: (i) a subset of patients with right-sided prostate cancer; (ii) a subset of patients with left-sided prostate cancer (Table 4); (iii) a subset of patients with both-sided prostate cancer. It is worthwhile noting that the RES of the right-side of the prostate was significantly lower than the left-side in the left- and both-sided cancer patients group. Moreover, PHASE of the left-side of the prostate was significantly lower than the right-side in the both-sided cancer patients’ group.

### Results from the multi-feature decision support system

Using the parameter set comprising BMI, PSA Density, PSA, AGE, RES, PHASE, and REA, we built a SVM multi-feature computational model as described above and derived cancer recognition accuracy, sensitivity, specificity, positive and negative predictive values (PPV and NPV).

The average prediction accuracy achieved is shown in Figure 4, with a final score as of 75.00%. (**Fig.2**). This was obtained mathematically combining the following four parameters that were identified as clinically relevant: BMI, PSA Density, RES, and PSA. A comprehensive, ranked clinical feature list for this decision support system is reported in Table 6, while the corresponding confusion matrix is in Table 7. Sensitivity and specificity of the PCa prediction vs. BPH were 63.33% and 83.75% respectively. The PPV and NPV were 74.51% and 75.28%, respectively. It is worthwhile noting that the resistance (averaged between the right and left prostate lobes) is one of the most informative features and gives a significant contribution to achieve the 75.00% of accuracy.

**Figure 2:**
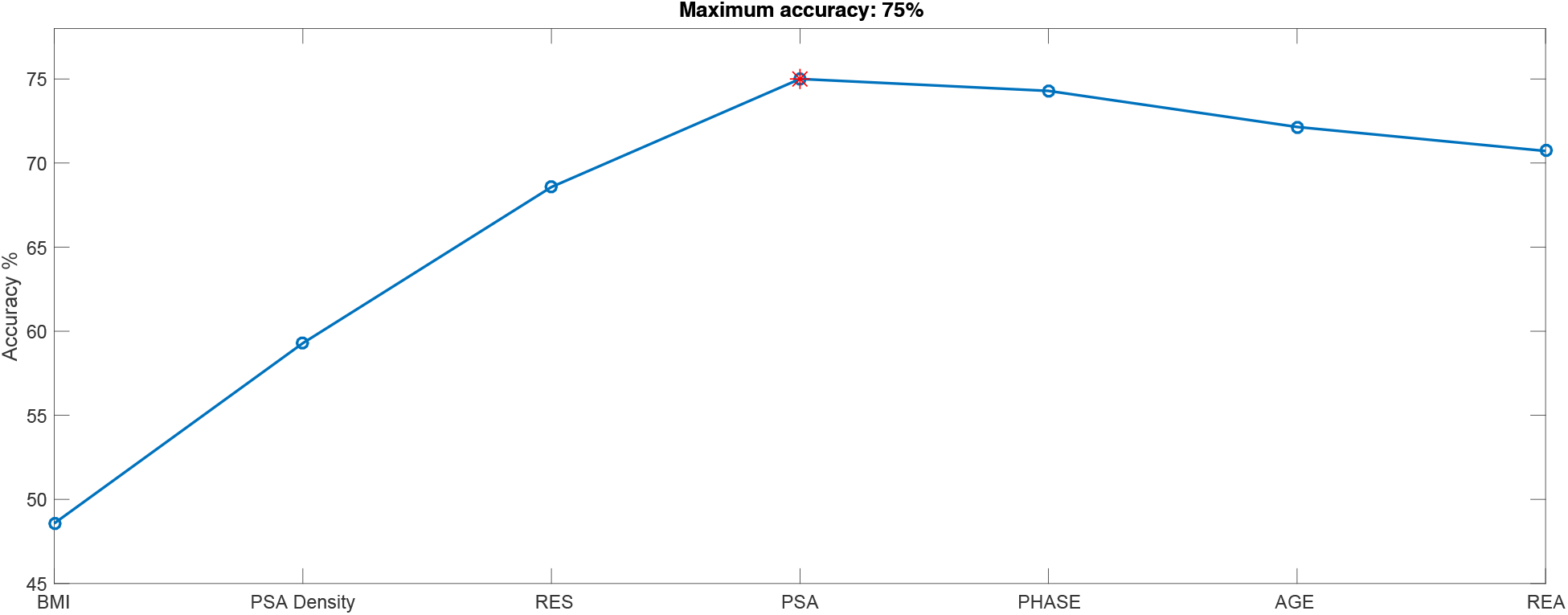
Accuracy trend of Cancer vs BPH recognition problem as a function of incremental number of features. The features are ranked according to the SVM-RFE criterion

Importantly, as a counterproof, we obtained a significant decrease in the PCa prediction accuracy of 62.86% (see details in Table 8) while repeating the same SVM-based computational procedure using a feature-set that does not include the three biometric measures. As expected, the most informative sub-features set included BMI, PSA Density, and PSA.

## Discussion

Bioelectric Impedance Activity (BIA) of different tissues was originally investigated by Geddes and Baker in the ‘60s [18]. They carried out an electrical measurement on living tissues demonstrating different values of resistivity. From that period, the BIA test have been used for various purposes such as the lean and fat body mass calculation and other medical applications like skin and breast cancer diagnosis. [19–21]. Halter et al. measured electrical properties of “ex vivo” prostate tissues with the aim of future applications for PCa non-invasive diagnosis. They realized that PCa, BPH, non-hyperplastic glandular tissue and stromal tissue had different conductivity at all frequencies while mean cancer permittivity was significantly greater than that of benign tissues at high frequencies. [12]. Other authors demonstrated that best results for cancer diagnosis by BIA test were obtained by measuring the tissue phase angle. Low phase angle suggests cell death or decreased cell integrity, whereas higher phase angle suggests healthy cell [22,23]. A low phase angle has been associated with an impaired outcome in tumor diseases such as pancreatic cancer, colorectal cancer and lung cancer [5,6,9]. Tyagi et al recently demonstrated that low phase angle values measured by BIA test allow for discriminating PCa patients from matched controls and those with advanced stage and high risk PCa in particular. They investigated a group of subjects using the BIA electrodes placement on the right upper and right lower limb. On the other hand, all PCa diagnosed subjects had a total PSA increased values and other concomitant diseases excluded to avoid the risk of false positive results [11]. Similarly, Khan developed a new composite impedance metrics method with a 9-electrode micro endoscopic probe. This novel device was tested on “in vivo” and “ex vivo” prostate tissue either intra-operatively or after the prostate removal in patients who underwent to surgery for PCa or BPH. The results obtained demonstrated a predictive accuracy of 90.79% for PCa [7]. For these reasons, we provided an alternative electrode placement and a restricted loco-regional electric field in order to improve the BIA test sensitivity and specificity and reduce possible confounding factors. The finger probe allows the obtainment of the tissue resistance, reactance and phase angle measurements directly from the prostate gland surface through a restricted electric field generated into the pelvic bone girdle. However, prostate tissue presents an extreme variability of electrical absorption due to its water and/or stromal content and the presence of micro-calcifications in its tissue context with subsequent false positive results. Our results demonstrated that the finger probe is a promising, reliable, and easy-to-use tool to improve the accuracy of PCa non-invasive diagnosis together other standard clinical parameters. In patients where PCa was diagnosed in both prostate lobes (i.e., 65% of cases), BIA phase angles were found significantly different between the right- and left-side, whilst seemed to be comparable when PCa was diagnosed in a single lobe. For this reason, measurements from the BIA test were averaged between the two lobes to avoid inhomogeneous results within the cancer group patients for the development of the multi-feature decision support system. Our experimental evidences on BIA phase angles do not replicate previous findings reported in [8]. This may be justified by the presence of more represented stromal tissue and/or calcifications inside the gland, as well as by the normalization procedure that we have performed prior to the statistical analyses. All BIA measures including RES, REA, and PHASE, in fact, were normalized by dividing their value by the prostate volume estimated during the TRUS examination to avoid biases. Without normalization, patients with BPH and with PCa vs. healthy controls showed significant differences in terms of BIA phase angle (p=0.006 and 0.003, respectively), therefore confirming previous observations by Tyagi et al. [11]. BIA resistance values were lower in patients with PCa although, taken alone, it seemed to be unable to differentiate cancer from non-cancer patients, whilst it was significantly different between healthy subjects and the BPH group. BIA reactance values were significantly different between healthy subjects and patients, although taken alone were not significantly different between BPH and PCa patients.

In this sense, likewise for the PSA alone, the BIA test failed to differentiate subjects with clinical suspicion of PCa and prospectively missed the intent of avoiding unnecessary biopsies. Nevertheless, when combined with the other standard clinical parameters including patients’ PSA and PSA density, BIA test provided meaningful information for discerning between PCa and BPH patients with an accuracy as high as 75% at a single patient level.

Our results indicates a good PCa prediction using a combination of the following clinical features: BMI, Age, PSA, PSA Density. In this case, sensitivity and specificity are lower than the ones associated with a combination of BMI, PSA Density, RES, and PSA, thus demonstrating the significant clinical information associated with BIA test.

### Limitations of the study

Study limitations include the small amount of data, especially from healthy volunteers.

## Conclusions

The use of a novel transrectal “finger-probe” allows to perform the BIA test with a minimal discomfort for the patient, contributing to an accuracy as high as 75% for PCa vs. BPH prediction when properly combined with BMI, total PSA, and PSA density. Interestingly, the test can be easily repeatable. Hence, the BIA test may be useful to decrease the need of useless biopsies.

